# FBXO45-MYCBP2 regulates mitotic cell fate by targeting FBXW7 for degradation

**DOI:** 10.1101/493791

**Authors:** Kai T. Richter, Yvonne T. Kschonsak, Barbara Vodicska, Ingrid Hoffmann

## Abstract

Cell fate decision upon prolonged mitotic arrest induced by microtubule targeting agents depends on the activity of the tumor suppressor and F-box protein FBXW7. FBXW7 promotes mitotic cell death and prevents premature escape from mitosis through mitotic slippage. Mitotic slippage is a process that can cause chemoresistance and tumor relapse. Therefore, understanding the mechanisms that regulate the balance between mitotic cell death and mitotic slippage is an important task. Here we report that FBXW7 protein levels markedly decline during extended mitotic arrest. FBXO45 binds to a conserved acidic N-terminal motif of FBXW7 specifically under a prolonged delay in mitosis, leading to ubiquitylation and subsequent proteasomal degradation of FBXW7 by the FBXO45-MYCBP2 E3 ubiquitin ligase. Moreover, we find that FBXO45-MYCBP2 counteracts FBXW7 in that it promotes mitotic slippage and prevents cell death in mitosis. Targeting this interaction represents a promising strategy to prevent chemotherapy resistance.

## INTRODUCTION

The attachment of a single ubiquitin molecule or a polyubiquitin chain to a eukaryotic protein is an essential signaling event that can profoundly affect the fate of the target protein. For example, the ubiquitin-proteasome system controls protein degradation in a broad array of cellular processes [1]. Cancer cells often contain mutations targeting ubiquitin-mediated proteolysis that lead to tumorigenesis. FBXW7, an F-box protein and substrate receptor for the SCF (SKP1-CUL1-F-box protein) E3 ubiquitin ligase complex, acts as a tumor suppressor and is mutated or deleted in a variety of human cancers [2]. FBXW7 exerts its anti-tumor activity through destruction of key oncoproteins, including JUN [3] [4], MYC [5] [6], Cyclin E [7] [8] [9] and Notch1 [10] [11].

The abundance of F-box proteins including FBXW7 is controlled by ubiquitylation and degradation in an autocatalytic reaction within the SCF complex [12]. FBXW7 autoubiquitylation can be regulated by Glomulin (GLMN), a protein that binds to RBX1 leading to inhibition of SCF activity, thereby increasing FBXW7 protein levels [13] [14]. The only ubiquitin ligase known to date involved in regulating FBXW7 levels in neurons is Parkin. However, this regulation has only been observed for the FBXW7β isoform [15]. It has also been shown that PLK2-kinase dependent phosphorylation of FBXW7 at S176 leads to its ubiquitin-mediated degradation and stabilization of Cyclin E [16].

Accurate spindle function is crucial for a successful mitosis. Perturbation of microtubule dynamics leads to sustained activation of the spindle assembly checkpoint (SAC) [17]. The SAC delays exit from mitosis by preventing the anaphase-promoting complex/cyclosome (APC/C) mediated proteolysis of Cyclin B1. Anti-cancer chemotherapeutics including vinca alkaloids or taxanes target microtubules and are successfully used in the clinics to treat multiple types of cancer. Anti-microtubule drugs cause an arrest or prolonged delay in mitosis followed by mitotic cell death. On the other hand, slow degradation of Cyclin B1 during a prolonged mitotic arrest can cause cells to prematurely exit from mitosis, a process called mitotic slippage [18]. During mitotic slippage, cells do not undergo proper chromosome segregation and cytokinesis. Most of the resulting tetraploid cells either undergo cell death after mitosis or arrest in interphase. However, depending on the p53 status of these cells, they may continue to proliferate as genomically unstable cells. This can lead to chemoresistance, thereby limiting the therapeutic use of anti-microtubule drugs [19] [20]. FBXW7 is a known regulator of mitotic cell fate. Upon mitotic arrest, FBXW7 promotes mitotic cell death and prevents mitotic slippage [21] [22].

FBXO45 (FSN-1 in *C. elegans*; DFsn in *Drosophila melanogaster*) is an evolutionary conserved F-box protein. FBXO45 is atypical in that it binds to SKP1 via its F-box motif, but recruits an alternate RING-finger protein known as MYCBP2, a member of the PHR (PAM/Highwire/RPM-1) protein family and E3 ubiquitin ligase. MYCBP2 was shown to have important functions in developmental processes, such as axon termination and synapse formation, as well as axon degeneration (reviewed by [23] [24]). FBXO45 contains a conserved F-box domain and a SPRY domain, which recruits substrates to the ubiquitin ligase complex [25] [26]. FBXO45-MYCBP2 has been linked to the proteasomal degradation of a few targets including the DLK-1 and p38 MAP kinase pathway [27] and the nicotinamide-nucleotide adenylyltransferase family member NMNAT [28].

Here we report the identification of the E3 ligase FBXO45-MYCBP2 as a regulator of FBXW7 abundance during prolonged mitotic arrest induced by spindle poisons. FBXO45 binds to a conserved acidic N-terminal region of FBXW7. Depletion of FBXO45 leads to stabilization of FBXW7 protein upon mitotic arrest. Ubiquitylation of FBXW7 by FBXO45-MYCBP2 induces its proteasomal degradation inducing an increase in mitotic slippage and prevention of mitotic cell death.

## RESULTS AND DISCUSSION

### Prolonged mitotic arrest leads to reduced FBXW7 protein levels

The SCF-FBXW7 complex acts as a key factor determining sensitivity to antimitotic drugs in cancer by inducing mitotic cell death and preventing mitotic slippage [29] [22] [21]. As mitotic slippage frequently occurs under prolonged mitotic arrest, it is conceivable that this could be induced by a decrease in FBXW7 protein levels. To address this, we analyzed protein levels of the ubiquitously expressed α-isoform of FBXW7 in mitotic HeLa cells at different time points after induction of mitotic arrest. We observed a slow but gradual decrease in the amount of FBXW7 protein during prolonged mitotic arrest (Figure 1A). The dynamics of this decrease was similar to the decay of two other regulators of mitotic cell fate, Cyclin B1 and MCL-1 [18] [30]. Moreover, the decay of FBXW7, Cyclin B1 and MCL-1 was prevented by the addition of the proteasomal inhibitor MG132 (Figure 1B). We therefore anticipated that FBXW7 protein levels could be decreased in response to proteasomal protein degradation. To identify proteins that regulate SCF-FBXW7 protein levels we made use of an unbiased screen where we aimed at identifying FBXW7 binding proteins. We expressed Flag-FBXW7α in HEK-293T cells, FBXW7α complexes were immunoprecipitated and subsequent mass spectrometrical analysis was performed. Among the identified peptides were sequences corresponding to FBXO45 and MYCBP2, along with SCF components and known substrates of FBXW7 including MYC, Notch1/2 and members of the mediator complex (Figure 1C, Table S1). Interestingly, FBXO45 and MYCBP2 have previously been identified in other screens for FBXW7 interacting proteins [31] [32]. They have also been described to form an SCF-like E3 ubiquitin ligase complex [33]. We therefore analyzed whether FBXO45-MYCBP2 is the ubiquitin ligase that regulates FBXW7 protein levels under prolonged mitotic arrest.

**Figure 1.**
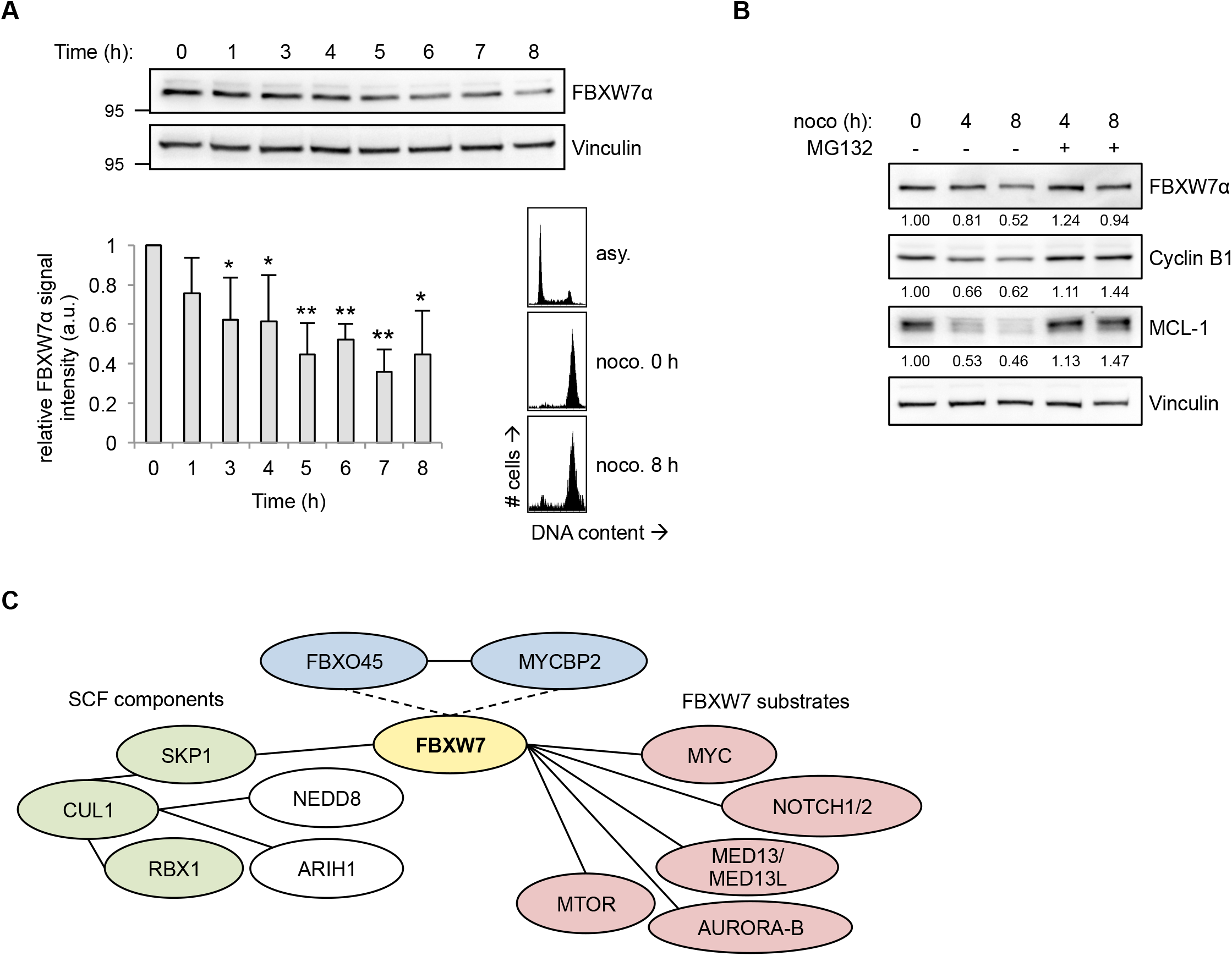
FBXW7 protein levels are reduced under prolonged mitotic arrest. (A) FBXW7α protein levels decrease during prolonged mitotic arrest. HeLa cells were treated with 250 ng/ml nocodazole for 2 h. Mitotic cells were collected by mitotic shake-off (time point 0) and further incubated with 250 ng/ml nocodazole. The cells were harvested at different time points by a mitotic shake-off. (Top) Cell extracts were prepared and analyzed by Western blotting. (Bottom, left) Relative FBXW7α signal intensities were quantified. Average signal intensities and standard deviations from n=4 experiments were calculated. Statistical significance (difference to time point 0) was analyzed by a two-tailed, unpaired t-test with unequal variance. * p<0.05; ** p<0.01. (Bottom, right) Mitotic arrest was confirmed for two time points (0 h and 8 h) by FACS analysis. Asynchronous (asy.) cells served as a control. (B) Decrease in FBXW7α protein levels during prolonged mitotic arrest depends on proteasomal activity. HeLa cells were treated with 250 ng/ml nocodazole for 2 h. Mitotic cells were collected by mitotic shake-off (time point 0) and further incubated with 250 ng/ml nocodazole. MG132 was added in order to inhibit the proteasome. Cells were harvested at different time points by a mitotic shake-off. Cell extracts were prepared and analyzed by Western blotting. Relative FBXW7α, Cyclin B1 and MCL-1 signal intensities were quantified. (C) Network of known and putative FBXW7 interaction partners. Immunoprecipitation of Flag-FBXW7α from HEK-293T cells treated with the proteasomal inhibitor MG132 for 4 h and subsequent mass spectrometry analysis identified several known SCF components and the SCF regulators NEDD8 and ARIH1 as well as FBXW7 substrates. Moreover, FBXO45 and MYCBP2 were identified as FBXW7 interaction partners and could act as putative FBXW7 regulators.

### The FBXO45-MYCBP2 complex binds to a conserved N-terminal motif in FBXW7α

First, we checked whether FBXO45 and FBXW7α bind to each other and found that Flag-FBXO45 was able to co-immunoprecipitate endogenous FBXW7α (Figure 2A) and vice versa Flag-FBXW7α co-immunoprecipitated endogenous FBXO45 (Figure 2B). To confirm the specificity of the binding between FBXW7α and FBXO45, several other F-box and DCAF substrate receptors were expressed and immunoprecipitated. No interaction was detected between FBXO45/FBXW7α and the control proteins suggesting that the binding of FBXO45 and FBXW7α was specific (Figures 2A-B). FBXW7 occurs in three isoforms, FBXW7α, FBXW7β and FBXW7γ, that differ in subcellular localization and their N-terminal domain. FBXW7α is thought to perform most FBXW7 functions [2]. To identify the binding site of FBXO45 within FBXW7 we generated different FBXW7α truncated versions and found that aa106-126 in the N-terminal domain of FBXW7α were required for FBXO45 binding (Figures EV1A-D). Co-immunoprecipitation experiments using full-length Flag-FBXW7α or a Flag-FBXW7αΔ106-126 mutant showed that FBXW7αΔ106-126 failed to bind FBXO45 and MYCBP2 (Figure 2C). This could be confirmed by the fact that FBXO45 only interacted with FBXW7α but not with the other two isoforms, FBXW7β or FBXW7γ (Figure EV1E), that lack the FBXW7α-specific N-terminal domain [34] again suggesting FBXW7α specifically binds FBXO45. Interestingly, the N-terminal stretch within the FBXW7α isoform (aa106-126) harbors conserved acidic amino acid residues as a potential FBXO45 interaction domain (Figure 2D). On the other hand, we identified the central domain of MYCBP2 (aa1951-2950) to be responsible for FBXW7α binding (Figures EV2A-B). As this domain also contains the FBXO45 interaction site of MYCBP2, we hypothesized that FBXO45 could be the direct interaction partner of FBXW7α within the FBXO45-MYCBP2 complex. Analysis of binding between purified MBP-tagged N-terminal domain of FBXW7α (MBP-FBXW7α-N167) and *in vitro* translated [^35^S]-FBXO45 or [^35^S]-MYCBP2(1951-2950) showed that FBXW7α directly binds to FBXO45 but not to MYCBP2 (Figure 2E). Furthermore, using sequential co-immunoprecipitation experiments we showed that FBXW7α exists in a complex with FBXO45 and MYCBP2 (Figure 2F).

**Figure 2.**
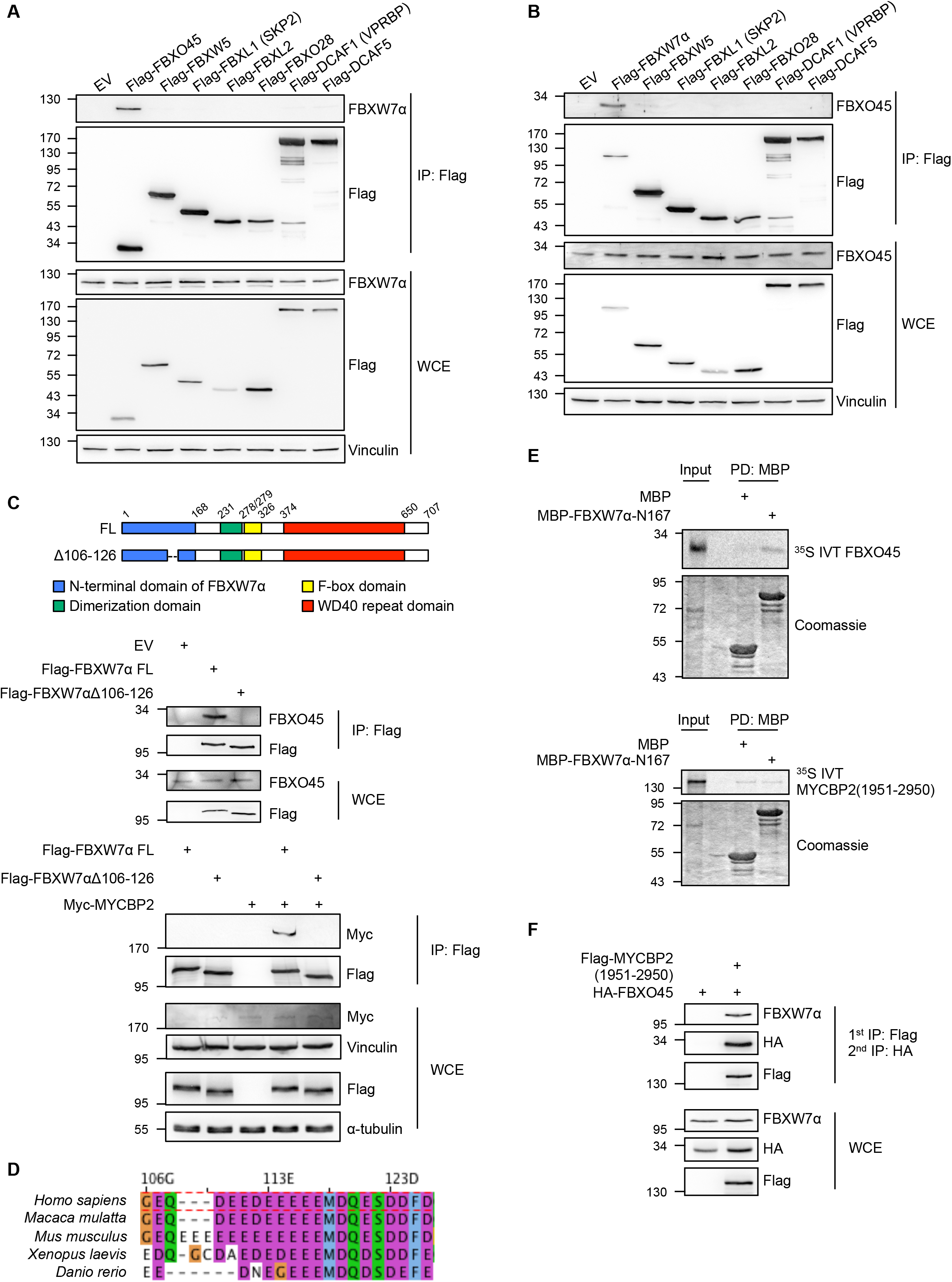
FBXO45-MYCBP2 binds to a conserved N-terminal motif of FBXW7α. (A-B) FBXO45 and FBXW7α specifically interact in reciprocal immunoprecipitations. Several Flag tagged F-box and DCAF proteins were overexpressed in HEK-293T cells for 24 h. Cell extracts were used for an immunoprecipitation directed against the Flag tag. (C) (Top) Schematic representation of Δ106-126 deletion in FBXW7α used in immunoprecipitation experiments. Positions of N-terminal domain (blue), dimerization domain (green), F-box domain (yellow) and WD40 domain (red) are indicated. (Bottom) FBXO45 and MYCBP2 interact with the N-terminal domain of FBXW7α. Flag-FBXW7α FL or Flag-FBXW7αΔ106-126 were overexpressed in HEK-293T cells for 24 h. Cell extracts were used for α-Flag immunoprecipitations. (D) Sequence alignment of amino acid residues 106-126 from human FBXW7α with homologues from *Macaca mulatta, Mus musculus, Xenopus laevis* and *Danio rerio*. (E) Either MBP alone or MBP-FBXW7α-N167 were incubated with *in vitro* translated and [^35^S]-methionine containing FBXO45 or MYCBP2(1951-2950). MBP was pulled down with amylose beads. Pull-down samples were analyzed by SDS-PAGE and Colloidal Coomassie staining. *In vitro* translated proteins were detected by autoradiography. (F) FBXW7α forms a complex with MYCBP2 and FBXO45. Flag-MYCBP2(1951-2950) and HA-FBXO45 were co-expressed in HEK-293T cells for 24 h. Cell extracts were used for an immunoprecipitation with α-Flag agarose beads. Immunoprecipitated proteins were eluted by competition with Flag peptide. Eluates were used for a second immunoprecipitation directed against the HA tag. Immunoprecipitates obtained after sequential immunoprecipitation were analyzed by Western blotting.

For some but not all F-box proteins it has been demonstrated that phosphorylation of their specific substrates is required for binding [35]. We have previously shown that phosphorylation of FBXW7 by PLK2 leads to destabilization of the F-box protein [16]. However, as shown in Figure EV2C, inhibition of PLK2 by the small molecule inhibitor BI2536 that targets both PLK1 and PLK2 kinases did not impair the binding between FBXO45 and FBXW7. Moreover, overexpression of GFP-PLK2 did not promote the interaction (Figure EV2D). Our data therefore suggest that the FBXW7α-FBXO45 interaction is independent of phosphorylation by PLK2.

Together, these results show that the FBXO45-MYCBP2 complex interacts with a conserved acidic stretch within the N-terminal domain of FBXW7α.

### FBXO45-MYCBP2 targets FBXW7α for degradation specifically during mitotic arrest

As FBXW7α is predominantly found in the nucleus [36] whereas FBXO45 and MYCBP2 are cytosolic proteins [25] [37] (Figure 3A), a possible interaction between these proteins may occur in mitosis upon nuclear envelope breakdown, when the contents of the cell nucleus are released into the cytoplasm. As FBXO45 is likely to be the substrate binding factor we analyzed the interaction between Flag-FBXO45 and endogenous FBXW7α in asynchronous and mitotic HeLa cells. We specifically prepared the cell extracts for the experiment shown in Figure 3B under conditions where the nucleus remains intact in asynchronous cells. Interestingly, FBXW7α was specifically found in co-immunoprecipitation with Flag-FBXO45 in mitotic cells (Figure 3B). Furthermore, we noticed an increase in FBXW7α protein levels upon siRNA-mediated downregulation of FBXO45. This effect could only be observed in cells under a prolonged mitotic arrest but not in cells passing through an unperturbed mitosis. In addition, the effect was not observed in asynchronous cells (Figures 3C and EV3A-B). On the other hand, FBXW7 downregulation had no effect on FBXO45 protein levels (Figure 3C). To exclude off-target effects we used different FBXO45 and MYCBP2 siRNAs to reduce their expression in HeLa cells and confirmed that FBXW7α protein levels were upregulated in cells that were FBXO45 or MYCBP2-depleted under prolonged mitotic arrest (Figure EV3C). The effect could be confirmed in U2OS cells using nocodazole and in HeLa cells using different inhibitors that cause a mitotic arrest (Figures EV3D-I). We also aimed to confirm the observed regulation of FBXW7α protein levels by expression of a dominant-negative MYCBP2(1951-2950) fragment that binds FBXO45 and FBXW7α but lacks the RING domain (Figures EV2A-B). Upon ectopical expression of Flag-MYCBP2(1951-2950) we found that FBXW7α protein levels were specifically upregulated in mitotic cells after treatment with nocodazole (Figure 3D). Taken together, FBXO45 binds FBXW7α in mitotic cells and the FBXO45-MYCBP2 complex regulates FBXW7α protein levels specifically during mitotic arrest.

**Figure 3.**
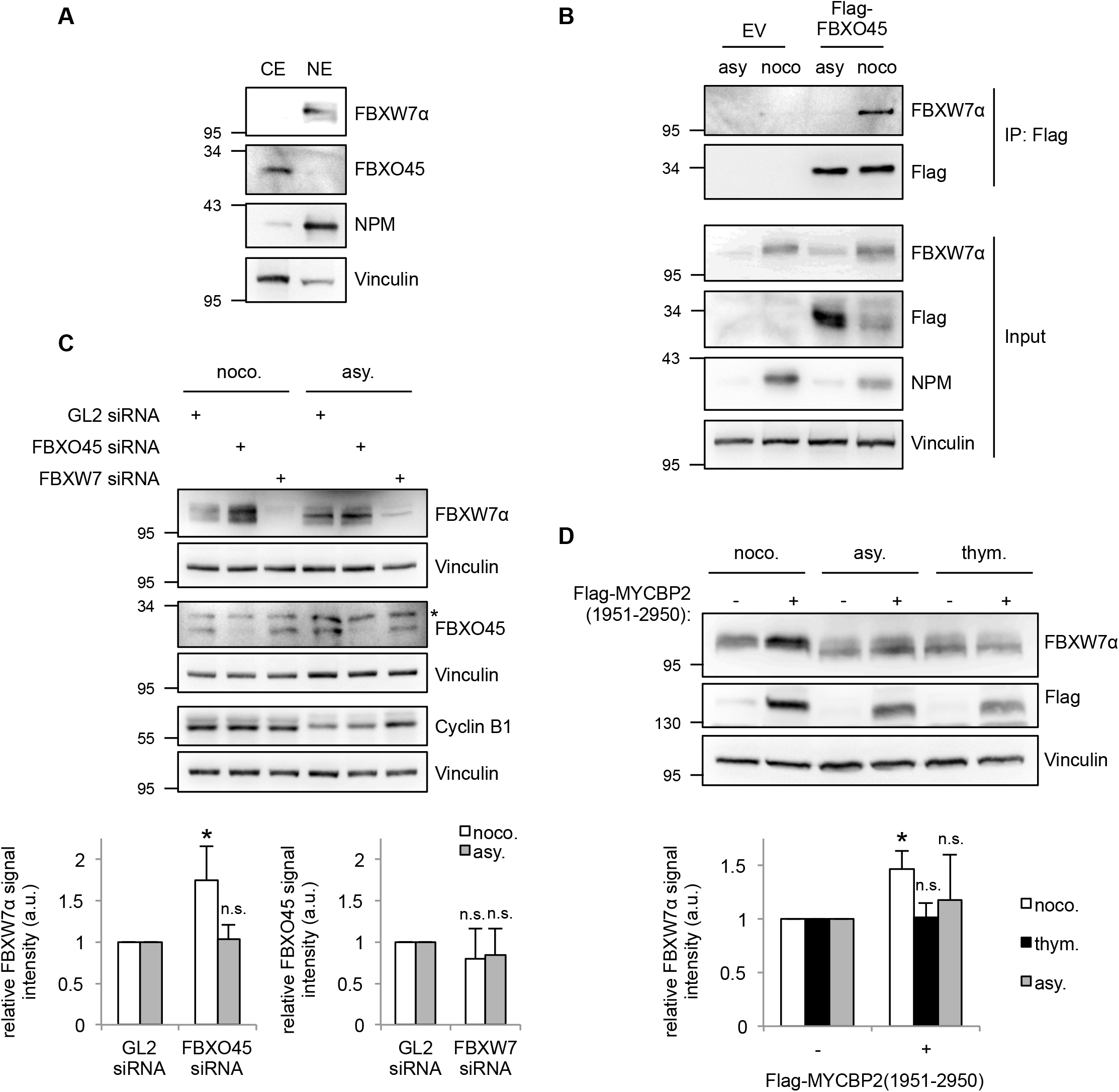
FBXO45 and MYCBP2 downregulate FBXW7α protein levels during mitotic arrest. (A) FBXW7α is a nuclear protein, whereas FBXO45 is localized in the cytoplasm. Cytoplasmic and nuclear extracts were prepared from HeLa cells and analyzed by Western blotting. (B) FBXO45 and FBXW7α interact predominantly in mitosis. Flag-FBXO45 was overexpressed in HeLa cells. Asynchronous or mitotic cells that had been treated with nocodazole for 17 h were harvested. Cytoplasmic extracts were used for an immunoprecipitation directed against the Flag tag. (C) HeLa cells were transfected with 30 nM FBXO45 or FBXW7 siRNAs for 72 h. GL2 siRNA was used as a control. The cells were arrested in mitosis by nocodazole (noco.) treatment for 17 h and collected by a mitotic shake-off. Mitotic cells were compared with an asynchronous (asy.) cell population. (Top) Cell extracts were analyzed by Western blotting. Crossreacting band in FBXO45 immunoblot is marked with an asterisk. As the samples were analyzed in three Western blots, three vinculin immunoblots are shown as loading controls. (Bottom) Quantification of relative FBXW7α and FBXO45 signal intensities are shown. Relative FBXW7α and FBXO45 signals in the GL2 controls were set to 1. Average signal intensities and standard deviations from n=4 experiments were calculated. Statistical significance was analyzed by a two-tailed, unpaired t-test with unequal variance. * p<0.05; n.s.: not significant. (D) Expression of Flag-MYCBP2(1951-2950) was induced in a stable U2OS Flp-In-T-Rex cell line by doxycycline treatment. The cells were arrested in mitosis by nocodazole treatment (noco.). Alternatively, they were treated with thymidine (thym.) or left untreated (asy.). U2OS cells that do not express the MYCBP2 fragment served as controls. (Top) Cell extracts were analyzed by Western blotting. (Bottom) Quantification of relative FBXW7α signal intensity was performed as described in (C).

### FBXO45-MYCBP2 targets FBXW7α for ubiquitylation and degradation

To find out whether the regulation of FBXW7α by FBXO45-MYCBP2 is mediated by ubiquitylation we carried out *in vivo* ubiquitylation assays. We found that both overexpression of FBXO45 and MYCBP2 promote ubiquitylation of FBXW7α (Figure 4A). Deletion of the FBXO45 binding site in FBXW7α reduced the ubiquitylation of FBXW7α markedly in *in vivo* ubiquitylation assays using FBXW7αΔ106-126 expression (Figure 4B). To study the effect of FBXO45 on FBXW7α protein stability, we depleted FBXO45 in nocodazole-treated cells and added cycloheximide to block translation. Cells were then harvested at different time points after cycloheximide addition. In FBXO45-depleted cells FBXW7α was markedly stabilized (Figure 4C) suggesting that FBXO45 promotes degradation of FBXW7α upon mitotic arrest. In addition, Flag-FBXW7αΔ106-126 was stabilized in comparison to full-length Flag-FBXW7α. The stability of Flag-FBXW7α was increased by MG132 treatment (Figure 4D). From these experiments, we conclude that FBXO45-MYCBP2 mediated degradation of FBXW7α during prolonged mitosis depends on the proteasome.

**Figure 4.**
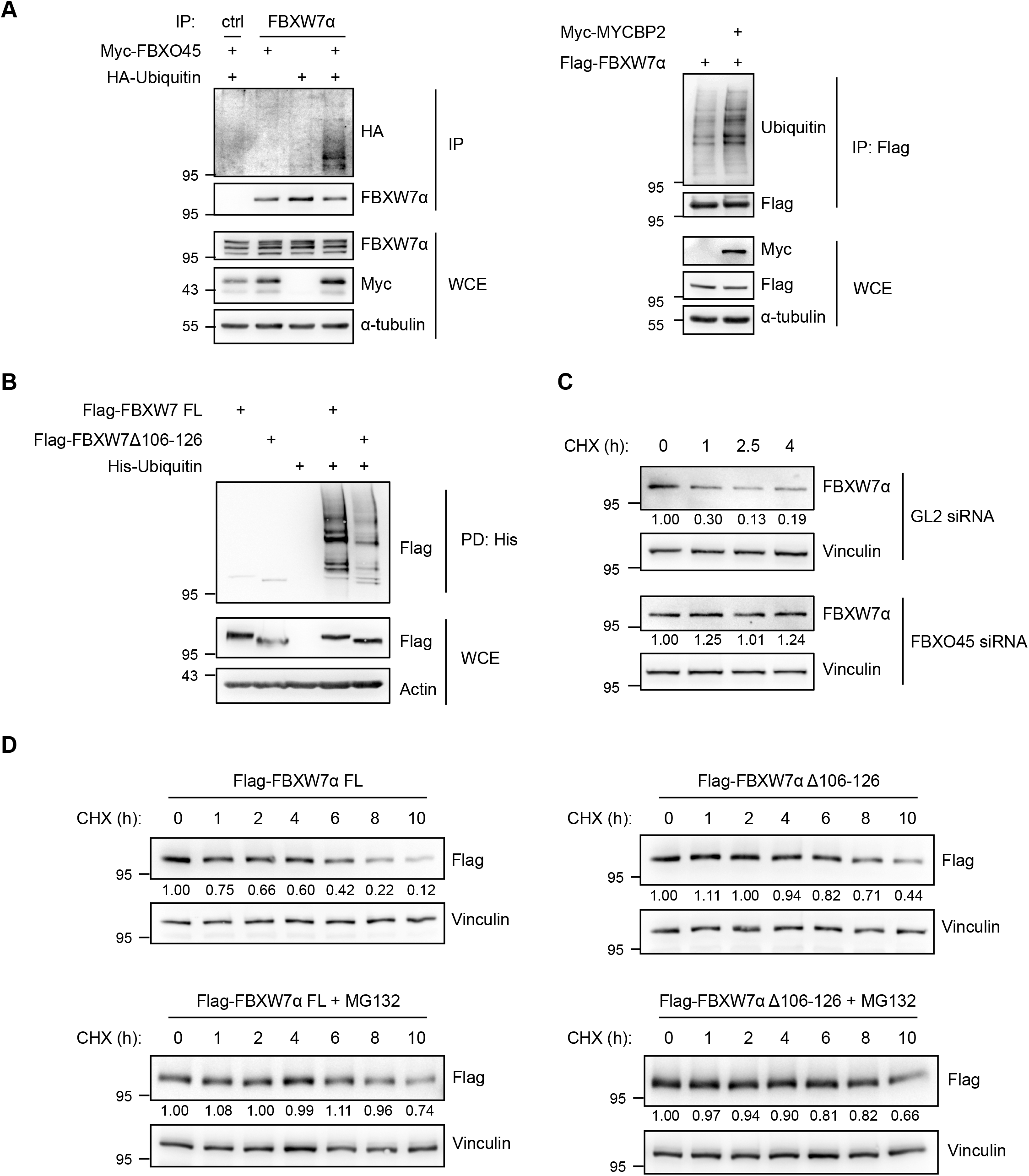
FBXO45 and MYCBP2 promote the ubiquitylation and destabilization of FBXW7α. (A) Myc-FBXO45 and HA-Ubiquitin or Myc-MYCBP2 and Flag-FBXW7α were overexpressed in HEK-293T cells for 48 h. Cells were arrested in mitosis by nocodazole treatment for 17 h. 4 h before harvesting, cells were treated with MG132. Cell extracts were used for immunoprecipitations directed against endogenous FBXW7α or against the Flag tag. (B) Flag-FBXW7α FL or Flag-FBXW7αΔ106-126 were co-expressed with His-Ubiquitin in HEK-293T cells. The cells were arrested in mitosis by nocodazole treatment. 4 h before harvesting, cells were treated with MG132. Cell extracts were prepared and α-His pull-downs were performed. (C) HeLa cells were transfected with 30 nM of GL2 or FBXO45 siRNA for 72 h. 17 h before harvesting, the cells were treated with nocodazole. Mitotic cells were collected by mitotic shake-off and further incubated with nocodazole and cycloheximide (CHX). The cells were harvested at different time points after the addition of cycloheximide by mitotic shake-off. Cell extracts were analyzed by Western blotting. Samples were analyzed on a single gel. Relative FBXW7α signal intensities were quantified. (D) Flag-FBXW7α FL or Flag-FBXW7αΔ106-126 were overexpressed in U2OS Flp-In-T-Rex cells by treatment with doxycycline. Cells were arrested in mitosis by nocodazole treatment. Cycloheximide (CHX) was added and cells were harvested at different time points after the addition of cycloheximide. MG132 was added in order to inhibit the proteasome. Cell extracts were analyzed by Western blotting. Relative Flag signal intensities were quantified.

### FBXO45-MYCBP2 reduces the sensitivity of cells to spindle poisons

Previous studies have shown that FBXW7 regulates mitotic cell fate. In fact, FBXW7 promotes mitotic cell death and prevents mitotic slippage in cells that have been treated with anti-microtubule drugs to arrest them in mitosis [21] [22], which is in line with its function as a tumor suppressor. As we had found that FBXO45-MYCBP2 regulates FBXW7α protein levels during mitotic arrest (Figure 3), we asked whether the FBXO45-MYCBP2 complex as a regulator of FBXW7 would also affect mitotic cell fate. To test this, we analyzed mitotic cell fate by live-cell imaging (Figure 5A, Video S1-2). For our live-cell imaging analysis, we treated the cells with spindle poisons at concentrations that completely prevented cell division and caused either mitotic cell death or mitotic slippage [38]. As shown in Figure 5B, reduction of FBXO45 led to an increase in mitotic cell death in cells that were arrested in mitosis with nocodazole. Similar effects were observed in cells arrested in mitosis by Taxol or vincristine treatment (Figures EV4A-B). To exclude off-target effects we used siRNAs targeting different regions of FBXO45 mRNA (Figure 5B). In addition, expression of an siRNA-resistant version of Flag-FBXO45 was able to rescue the effect on mitotic cell fate (Figure 5C). Moreover, siRNA-mediated downregulation of MYCBP2 also caused an increase in mitotic cell death (Figure EV4C). In contrast, FBXO45 and MYCBP2 depletion did not have an effect on progression of cells through an unperturbed mitosis (Figure EV4D). To verify the results in an siRNA independent approach, we analyzed the effect of the dominant-negative MYCBP2(1951-2950) fragment on mitotic cell fate. As expected, overexpression of Flag-MYCBP2(1951-2950) increased mitotic cell death (Figure 5D). Taken together, our results showed that the FBXO45-MYCBP2 complex negatively regulates mitotic cell death and increases mitotic slippage, suggesting that it has an opposing function during mitotic arrest compared to FBXW7. This is in line with the FBXO45-MYCBP2 complex being a negative regulator of FBXW7α during mitotic arrest. We therefore asked whether the observed effect of FBXO45-MYCBP2 on mitotic cell fate depends on FBXW7. In fact, FBXW7 depletion was able to rescue the effect of FBXO45, suggesting that FBXW7 is the downstream target of the FBXO45-MYCBP2 complex that mediates mitotic cell fate regulation (Figure 5E). Together, our data strongly suggest that FBXO45-MYCBP2-mediated degradation of FBXW7 under prolonged mitotic arrest induces mitotic slippage and prevents mitotic cell death.

**Figure 5.**
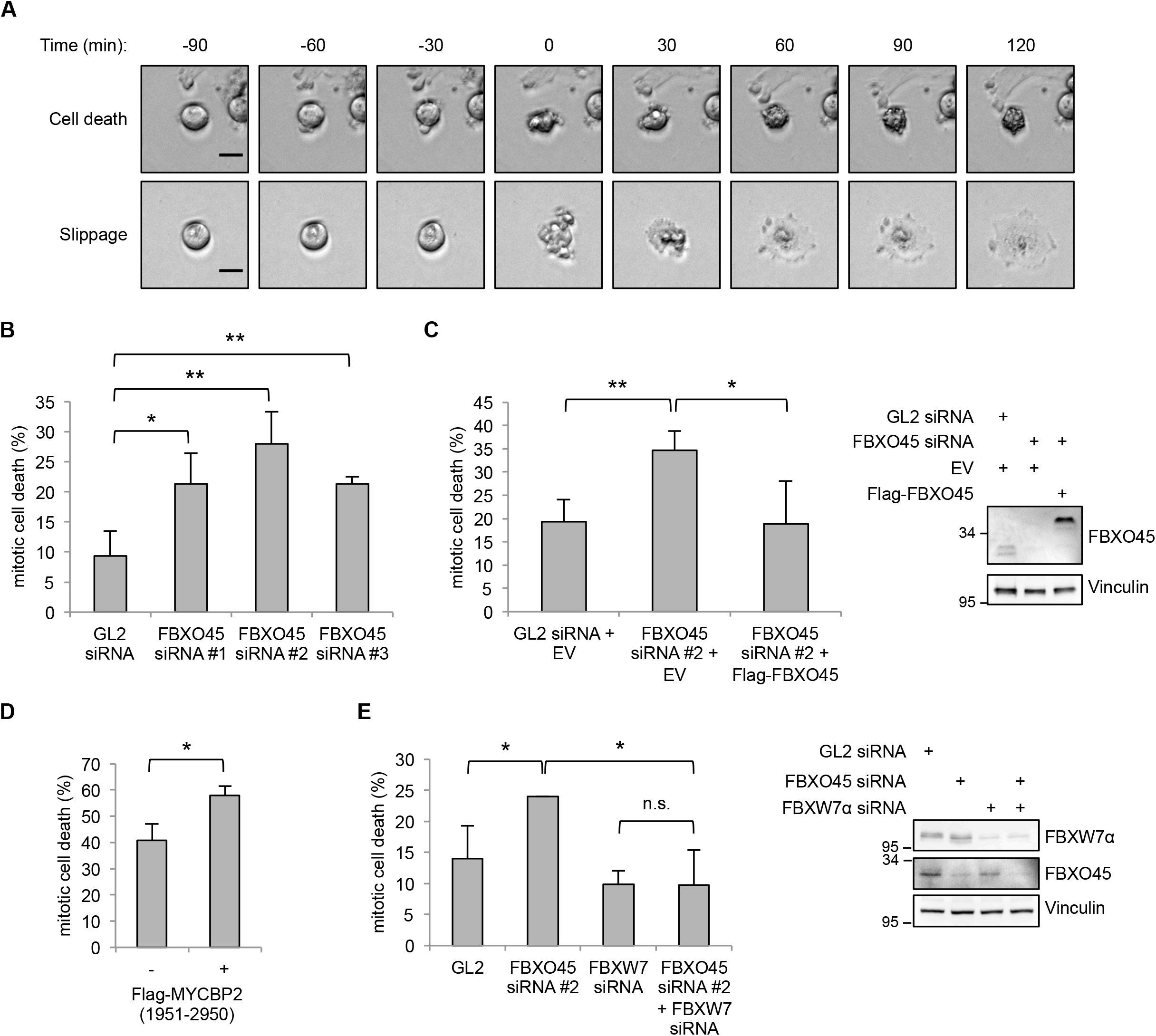
FBXO45 and MYCBP2 cause mitotic slippage. (A) U2OS cells were treated with 250 ng/mL nocodazole. 4 h after nocodazole addition, cells were analyzed by live-cell imaging. Representative images from live-cell imaging are shown. Time point 0 marks induction of mitotic cell death or mitotic slippage. Scale bars: 20 μm. (B) U2OS cells were transfected with 30 nM of GL2 or different FBXO45 siRNAs for 48 h and further incubated with 250 ng/mL nocodazole. Percentages of cells undergoing mitotic cell death were quantified. Cells from n=3 independent experiments were analyzed. In each experiment, 50 cells were quantified. (C) U2OS cells were cotransfected with 40 nM of GL2 or FBXO45 siRNA and an empty vector (EV) or an siRNA resistant version of Flag-FBXO45. 48 h after transfection, the cells were treated with nocodazole and mitotic cell fate was analyzed by live-cell imaging. Cells from n=4 independent experiments were quantified. (D) Expression of Flag-MYCBP2(1951-2950) was induced in U2OS cells by treatment with doxycycline for 48 h. Cells were arrested in mitosis by treatment with nocodazole and mitotic cell fate was analyzed by live-cell imaging. Cells from n=3 independent experiments were analyzed. (E) U2OS cells were transfected with 30 nM of GL2, FBXO45 or FBXW7 siRNA or co-transfected with FBXO45 and FBXW7 siRNA. 48 h after transfection, the cells were treated with nocodazole and mitotic cell fate was analyzed by live-cell imaging. Cells from n=3 independent experiments were quantified. For statistical analysis in (B)-(E), significance was analyzed by a two-tailed, unpaired t-test. * p<0.05, ** p<0.01, n.s.: not significant.

Prolonged mitotic arrest is caused by a sustained activation of the spindle assembly checkpoint (SAC). If the SAC cannot be satisfied and mitosis cannot be completed, there are two alternative cell fates. Cells either undergo mitotic cell death or perform mitotic slippage. Cell fate following mitotic arrest is regulated by two competing networks, namely Cyclin B1 degradation and pro-apoptotic caspase activation [39]. Besides cell fate regulation through Cyclin B1, MCL-1, a pro-survival member of the BCL-2 family, was shown to be involved in cell fate decision. MCL-1 is degraded by a ubiquitin-proteasome-dependent mechanism in response to the disruption of mitosis, resulting in cell death [39] [30] [22].

The tumor suppressor FBXW7 also plays important roles in mitotic cell fate decision [22] [29] [21]. However, whether or not FBXW7 has a direct role in regulating MCL-1 levels is a matter of debate [22] [40] [29] and therefore the identification of the substrate(s) of FBXW7 involved in regulating mitotic cell fate is an important task for future studies.

Our work identifies FBXO45 as a protein binding to a conserved acidic amino acid motif (aa106-126) of FBXW7α (Figures 1–2) to form a ternary complex with MYCBP2. Interestingly, available TCGA data show that FBXO45 is frequently overexpressed in different cancer types (cBioPortal for Cancer Genomics) [41] [42]. Therefore, it could be meaningful to further characterize FBXO45 as a putative oncoprotein in future studies. MYCBP2 has been recently uncovered as a novel type of E3 ligase with esterification activity leading to ubiquitylation of serine and threonine residues in substrates [43], which is intrinsic to higher eukaryotes. In the future, it would therefore be interesting to see whether FBXW7 could be ubiquitylated on hydroxyl groups by MYCBP2.

As FBXW7α is a nuclear protein but FBXO45 is found in the cytoplasm (Figure 3A), the two proteins can only interact upon nuclear envelope breakdown (Figure 3B). However, since mitosis is a relatively short process in the eukaryotic cell cycle, the successful formation of a sustained interaction between FBXO45 and FBXW7α as well as FBXW7α ubiquitylation may require a longer arrest in mitosis. Interestingly, the degradation of two other regulators of mitotic cell fate, MCL-1 and Cyclin B1, has also been shown to occur in a slow and gradual manner during mitotic arrest [18] [30].

In the future, the development of a specific inhibitor to block the FBXO45-MYCBP2-FBXW7α axis might actively promote mitotic cell death and prevent mitotic slippage. Therefore, treatment of cancer patients with this inhibitor after or in combination with anti-microtubule drugs could be a promising approach in order to increase the efficiency of chemotherapeutic treatment.

## MATERIALS AND METHODS

### Cell culture and transfection

Human cell lines were grown in a humidified atmosphere with 5% CO_2_ at 37°C. HeLa, U2OS and U2OS Flp-In-T-Rex cells were cultured in Dulbecco’s Modified Eagle’s Medium (DMEM, Sigma-Aldrich) with 1 g/l glucose. HEK-293T cells were grown in DMEM with 4.5 g/l glucose. The media were supplemented with 10% fetal bovine serum (FBS, Sigma-Aldrich or Thermo Scientific). HeLa GFP-α-tubulin/RFP-H2B cells (D. Gerlich, ETH Zürich) were cultured in DMEM containing 1 g/l glucose, 10% FBS, 500 μg/ml G418 and 0.5 μg/ml puromycin. The T-Rex system was used to generate U2OS Flp-In-T-Rex Flag-FBXW7α, U2OS Flp-In-T-Rex Flag-FBXW7αΔ106-126 and U2OS Flp-In-T-Rex Flag-MYCBP2(1951-2950) stable cell lines according to the manufacturer’s instructions (Life Technologies).

Cell culture work was performed under sterile conditions. Cell line authentication was regularly performed by Multiplexion in Heidelberg.

HEK-293T and HeLa cells were transiently transfected with plasmid DNA using polyethylenimine (PEI, Polysciences) at a final concentration of 5 μg/ml. The cells were incubated at 37°C and harvested 24-48 h after transfection. If the cells were harvested 48 h after transfection, the transfection mixture was removed 6-24 h after transfection and replaced by fresh growth medium.

For the transfection of HeLa or U2OS cells with siRNA, the transfection reagent Lipofectamine 2000 (Invitrogen) was used according to the manufacturer’s instructions. Cells were transfected with 30 nM of siRNA for 72 h. For rescue experiments, 50 ng of plasmid were co-transfected with 40 nM of siRNA for 72 h.

The following siRNA sequences were used:

> GL2 (firefly luciferase, control), 5’-CGUACGCGGAAUACUUCGAtt-3’;
>
> FBXO45 #1, 5’-GGAGAAAGAAUUCGAGUCAtt-3’;
>
> FBXO45 #2, 5’-ACACAUGGUUAUUGCGUAUtt-3’;
>
> FBXW7, 5’-ACAGGACAGUGUUUACAAAtt-3’;
>
> MYCBP2 #1, 5’-CCCGAGAUCUUGGGAAUAAtt-3’.

FBXO45 #3 (200933, L-023542-01-0005) and MYCBP2 #2 (23077, L-006951-00-0005) were purchased as SMARTpools from Dharmacon. If not stated otherwise, FBXO45 siRNA #2 was used for FBXO45 depletion and MYCBP2 siRNA #2 was transfected in order to downregulate MYCBP2.

### Cell cycle synchronization

In order to arrest HeLa or U2OS cells in G1/S phase, the cells were treated with 2 mM thymidine for 18 h. In order to arrest HeLa or U2OS cells in mitosis, cells were first synchronized in G1/S phase by treatment with 2 mM thymidine for 17 h. The cells were released into the cell cycle by washing three times with PBS and incubation with fresh growth medium for 5 h. Finally, the cells were treated with 250 ng/ml nocodazole, 500-1000 nM Taxol, 500 nM vincristine, 5 μM STLC or 100 nM BI2536 for 16-17 h. Mitotic cells were collected by a mitotic shake-off.

### MG132 and cycloheximide treatment

In order to inhibit proteasomal degradation in HEK-293T cells, the cells were treated with 10 μM MG132 (Sigma-Aldrich) for 4-5 h before harvesting. In order to inhibit protein synthesis in HeLa cells, they were treated with 100 μg/ml cycloheximide (Sigma-Aldrich). For a cycloheximide chase assay, the cells were harvested at different time points after cycloheximide treatment.

### Live-cell imaging

For the analysis of mitotic cell fate or unperturbed mitotic progression, live-cell imaging was performed. For mitotic cell fate analysis, U2OS cells were transfected with 30 nM siRNA targeting GL2, FBXW7, FBXO45 or MYCBP2. 48 h after transfection, 2.5 x 10^4^ cells were seeded into Ibidi dish chambers. 72 h after transfection, the cells were treated with 250 ng/ml nocodazole, 1 μM Taxol or 500 nM vincristine. 4 h after the spindle poison treatment, the cells were monitored by a 10x/0.3 EC PlnN Ph1 DICI objective on a Zeiss Cell Observer Z1 inverted microscope (AxioCam MRm camera system) with incubation at 5% CO_2_ and 37°C. Multi-tile phase-contrast images were taken every 10 min for 48 h using the Zeiss ZEN blue software. Data analysis was performed with ImageJ Fiji software. Cell death was defined by cell morphology and cessation of movement. Mitotic slippage was defined by mitotic exit without cell division. For the analysis of unperturbed mitotic progression, HeLa GFP-α-tubulin/RFP-H2B cells were transfected with 30 nM siRNA targeting GL2 or FBXO45 and MYCBP2. Analysis was performed as described previously [44]. 48 h after transfection, 2.5 x 10^4^ cells were seeded into Ibidi dish chambers and arrested in G1/S phase by thymidine treatment. 24 h after thymidine addition, the cells were released from the arrest and analyzed by live-cell imaging using LED module Colibri.2 with 470 nm for GFP and 590 nm for RFP fluorochrome excitation.

### Preparation of protein extracts from mammalian cells

For the preparation of cell extracts, cell pellets were resuspended in 3-5 volumes of RIPA, NP40 or Urea lysis buffer. RIPA lysis buffer (50 mM Tris-HCl pH 7.4, 1% NP40, 0.5% Na-deoxycholate, 0.1% SDS, 150 mM NaCl, 2 mM EDTA, 50 mM NaF, 1 mM DTT, 10 μg/ml TPCK, 5 μg/ml TLCK, 0.1 mM Na3VO4, 1 μg/ml Aprotinin, 1 μg/ml Leupeptin, 10 μg/ml Trypsin inhibitor from soybean) was used for the analysis of protein levels in whole cell extracts and for *in vivo* ubiquitylation assays. NP40 lysis buffer (40 mM Tris-HCl pH 7.5, 150 mM NaCl, 5 mM EDTA, 10 mM β-glycerophosphate, 5 mM NaF, 0.5% NP40, 1 mM DTT, 10 μg/ml TPCK, 5 μg/ml TLCK, 0.1 mM Na3VO4, 1 μg/ml Aprotinin, 1 μg/ml Leupeptin, 10 μg/ml Trypsin inhibitor from soybean) was used for immunoprecipitations. Urea lysis buffer (8 M Urea, 30 mM imidazole, 0.1 M phosphate buffer pH 8.0) was used for *in vivo* ubiquitylation assays. The cell lysates were incubated on ice for 30 min with short vortexing every 5-10 min. The lysates were cleared by centrifugation at maximal speed and 4°C in an Eppendorf 5415 R centrifuge for 15 min. The supernatants were transferred to fresh reaction tubes. Extracts of cytoplasmic and nuclear cell fractions were prepared using the CelLytic NuCLEAR Extraction Kit (Sigma) according to the manufacturer’s instructions.

The protein concentration of a cell extract was determined according to the Bradford method in a Bio-Rad Protein assay. For SDS-PAGE, the cell extracts were mixed with equal volumes of 2x Laemmli buffer and incubated at 95°C for 5 min.

### Western blotting

The following antibodies were used for Western blotting: Rabbit α-FBXW7α antibody (A301-720A, 1:15000) was obtained from Bethyl. Rabbit α-FBXO45 (NBP1-91891, 1:500) was purchased from Novus. Rabbit α-MYCBP2 was a kind gift from K. Scholich [45]. Mouse α-Flag (M2, F3165, 1:5000), mouse α-tubulin (B-5-1-2, 1:10000) and mouse α-Vinculin (hVIN-1, 1:5000) were obtained from Sigma-Aldrich. Rabbit α-Cyclin B1 has been described before [46]. Mouse α-HA (16B12, 1:1000) was purchased from Babco. Mouse α-Myc (9E10, 1:500), rabbit α-SKP1 (H-163, 1:1000) and mouse α-ubiquitin (P4D1, 1:1000) were obtained from Santa Cruz. Mouse α-Nucleophosmin (NPM, 32-5200, 1:1000) was ordered from Zymed. Rabbit α-MCL-1 (4572, 1:1000) was obtained from Cell Signaling. Goat α-mouse IgG HRP (1:5000) was purchased from Novus. Donkey α-rabbit IgG HRP (1:5000) was obtained from Jackson Laboratories.

### Immunoprecipitation

Immunoprecipitations of Flag-tagged proteins were performed using α-Flag M2 affinity beads (Sigma-Aldrich). For each immunoprecipitation reaction, 10-40 μl of the α-Flag M2 affinity bead suspension was used. The beads were washed twice with TBS, once with glycine buffer (0.1 M glycine-HCl pH 3.5) and three times with TBS. Buffers and beads were kept on ice and centrifugations were performed at 5000 rpm and 4°C for 2 min in an Eppendorf 5415 R centrifuge. HEK-293T or HeLa cell extracts were prepared with NP40 lysis buffer and 6-15 mg of each extract were transferred to the prepared beads. Each reaction was filled up to a final volume of 1 ml using NP40 lysis buffer. The reactions were incubated overnight on a rotating wheel at 4°C. After the incubation, the beads were washed 3-5 times with NP40 lysis buffer. Immunoprecipitated proteins were eluted from the beads by competition with 100 μl of a 3x Flag peptide solution (100-500 ng/μl in NP40 lysis buffer). The elution was carried out on ice for 30 min with short vortexing every 5-10 min. After the elution, the beads were centrifuged at 5000 rpm and 4°C for 2 min. 90 μl of the supernatant were transferred to a fresh reaction tube and 30 μl of 4x Laemmli buffer were added. After incubation at 95°C for 2 min, 20-40% of the immunoprecipitation sample were analyzed by SDS-PAGE and Western blotting.

In order to analyze whether proteins exist in ternary complexes, sequential immunoprecipitations were performed. In the first step of the experiment, 15-20 mg of HEK-293T cell extracts were used for an immunoprecipitation directed against a Flag-tagged MYCBP2 fragment. The immunoprecipitation was performed as described above. Instead of mixing the eluate with Laemmli buffer, it was used for a second immunoprecipitation directed against HA-FBXO45. HA-FBXO45 was immunoprecipitated with 20 μl of α-HA agarose beads (Sigma-Aldrich) according to the manufacturer’s instructions. Finally, the beads were incubated with 30 μl of 2x Laemmli buffer at 95°C for 5 min. The supernatant was analyzed by SDS-PAGE and Western blotting.

### Mass spectrometry analysis

Immunoprecipitation samples were analyzed by SDS-PAGE and stained by Colloidal Coomassie. Analysis was performed by M. Schnölzer (DKFZ Protein Analysis Facility, Heidelberg). Briefly, whole gel lanes were cut into slices. Proteins were reduced and alkylated by incubation with 10 mM DTT in 40 mM NH_4_HCO_3_ for 1 h at 56°C in the dark and incubation with 55 mM iodoacetamide in 40 mM NH_4_HCO_3_ for 30 min at 25°C. After washing of the slices with H2O and 50% acetonitrile, they were dried with 100% acetonitrile. Proteins were digested in-gel with trypsin (0.17 μg in 10 μl 40 mM NH_4_HCO_3_, Promega) at 37°C over night. Tryptic peptides were extracted with 50% acetonitrile/0.1% TFA and 100% acetonitrile. Supernatants were lyophilized and redissolved in 0.1% TFA/5% hexafluoroisopropanol. Solutions were analyzed by nanoLC-ESI-MS/MS. Peptides were separated with a nanoAcquity UPLC system (Waters GmbH). Peptides were loaded on a C18 trap column with a particle size of 5 μm (Waters GmbH). Liquid chromatography was carried out on a BEH130 C18 column with a particle size of 1.7 μm (Waters GmbH). A 1 h gradient was applied for protein identification. The nanoUPLC system was connected to an LTQ Orbitrap XL mass spectrometer (Thermo Scientific). Data were acquired with the Xcalibur software 2.1 (Thermo Scientific). The SwissProt database (taxonomy human) was used for database searches with the MASCOT search engine (Matrix Science). Data from individual gel slices were merged. Peptide mass tolerance was 5 ppm, fragment mass tolerance was 0.4 Da and significance threshold was p<0.01.

### *In vivo* ubiquitylation assays

HEK-293T cells transiently transfected with the indicated plasmids were treated with 10 μM MG132 4-5 h before harvesting. Cells were harvested and cell extracts were prepared with RIPA lysis buffer containing 10 mM of the DUB inhibitor N-ethylmaleimide (NEM). 2-3 mg of protein extract were used for each sample. Flag-FBXW7α was immunoprecipitated with α-Flag affinity beads. Immunoprecipitation and washing steps were performed with RIPA lysis buffer containing 10 mM NEM. Alternatively, endogenous FBXW7α was immunoprecipitated by incubation of each protein extract with 1 μg of FBXW7α antibody on a rotating wheel at 4°C overnight. After the incubation with the antibody, the extracts were incubated with 15 μl of protein A sepharose beads (GE Healthcare) on a rotating wheel at 4°C for 1 h. The beads were washed four times with RIPA lysis buffer containing 10 mM NEM. Finally, the beads were incubated in 30 μl of 2x Laemmli buffer at 95°C for 5 min. 25 μl of the supernatant were analyzed by SDS-PAGE and Western blotting.

For denaturing ubiquitylation assays, cell extracts were prepared using Urea lysis buffer. Cell extracts were sonified, centrifuged and incubated with Nickel-NTA agarose beads on a rotating wheel at RT for 2 h. Afterwards, the beads were washed four times with Urea lysis buffer and proteins were finally eluted using 30 μl of 2x Laemmli buffer containing 200 mM imidazole. 25 μl of the supernatant were analyzed by SDS-PAGE and Western blotting.

### Expression and purification of MBP-tagged proteins

Chemically competent *E. coli* Rosetta (DE3) were transformed with pMAL-MBP-FBXW7α-N167. A single colony was used to inoculate 10 ml of LB medium containing 100 μg/ml ampicillin and 0.2% glucose. The culture was incubated over night at 30°C with constant shaking at 180 rpm. The overnight culture was used to inoculate 1 l of LB medium containing 100 μg/ml ampicillin and 0.2% glucose. The culture was incubated at 37°C with constant shaking at 180 rpm. When an OD_600_ of 0.5 was reached, the culture was cooled down on ice at 4°C and protein expression was induced by the addition of 0.4 mM isopropyl-β-D-thiogalactopyranoside (IPTG). The culture was further incubated over night at 18°C with constant shaking at 180 rpm. Bacteria were harvested by centrifugation at 5000 rpm and 4°C for 15 min (F10-6x500Y rotor, Piramoon). The pellet was resuspended in 25 ml of cold column buffer. Cell lysis was performed with a high pressure homogenizer (15000-17000 psi/1030-1170 bar for one pass, EmulsiFlex C5, Avestin). The lysate was centrifuged at 20000 g and 4°C for 20 min (WX Ultra 80, Thermo Scientific). The supernatant was applied onto a column with 1 ml of equilibrated amylose resin (NEB). The column with the beads and the extract was incubated on a rotating wheel at 4°C for 1 h. Afterwards, the beads were washed three times with 15 ml of column buffer. Finally, the proteins bound to the beads were eluted with 5 ml of column buffer containing 10 mM maltose. 10 fractions of 500 μl each were collected. Protein containing fractions were identified by spotting the fractions on nitrocellulose and staining with Ponceau S solution. Protein containing fractions were pooled and MBP-FBXW7α-N167 was further purified with a preparative Superdex 200 column in 50 mM Tris-HCl pH 8.0, 100 mM NaCl, 5 mM β-mercaptoethanol, 5% glycerol. Purified MBP-FBXW7α-N167 fractions were analyzed by SDS-PAGE and Colloidal Coomassie staining. Protein containing fractions were aliquoted, frozen in liquid nitrogen and stored at −80°C.

### *In vitro* transcription and translation and *in vitro* binding assays

For *in vitro* binding assays, FBXO45 and MYCBP2(1951-2950) cDNA sequences in pCMV-3Tag1A backbones were transcribed and translated *in vitro* using the TNT T3 coupled reticulocyte system (Promega) according to the manufacturer’s instructions. The proteins were synthesized in the presence of 20 μCi [^35^S]-methionine so that synthesized proteins were radioactively labelled. 20 μl of the *in vitro* translated proteins were incubated with 10 μg of MBP-FBXW7α-N167 or MBP alone coupled to 10 μl of amylose beads in a final volume of 500 μl NP40 lysis buffer on a rotating wheel at 4°C for 2 h. The beads were washed five times with 800 μl of NP40 lysis buffer. Finally, the beads were incubated with 30 μl of 2x Laemmli buffer at 95°C for 5 min. Input and pull-down samples were analyzed by SDS-PAGE and stained with Colloidal Coomassie. The gel was then incubated with Amersham Amplify Fluorographic reagent (GE Healthcare) for 30 min with gentle shaking. Afterwards, the gel was dried at 80°C for 1 h in a vacuum dryer and analyzed by autoradiography.

### FACS analysis

For the analysis of mitotic arrest, the cellular DNA content was measured by FACS analysis. Cells were fixed with 70% ethanol at −20 °C and rehydrated in PBS for 15 min at RT. Afterwards, the cells were resuspended in 300 μl of PI staining solution (30 μg/ml propidium iodide, 10 μg/ml RNase A in PBS) and incubated for 15 min at RT. Stained cells were analyzed on a FACSCalibur (BD).

## ACKNOWLEDGMENTS

We thank D. Gerlich, L. Hengst, F. Melchior, G. Melino, A. Peschiaroli and K. Scholich for providing reagents and cell lines. We acknowledge the DKFZ Mass Spectrometry and Microscopy Core Facilities for providing equipment and excellent technical assistance. We thank Bettina Dörr for expert technical assistance. We acknowledge the members of our lab for critically reading the manuscript. This work was supported by a grant from the German Cancer Aid (Deutsche Krebshilfe) No. 110924 to I.Hoffmann.

## AUTHOR CONTRIBUTION

IH and KTR designed the experiments. KTR, YTK and BV performed the experiments. IH and KTR wrote the paper.

---

**Table S1.** Identification of FBXW7α interaction partners. Selection of proteins that were specifically identified in co-immunoprecipitation with Flag-FBXW7α. For each protein, the MASCOT score, mass, number of significant sequences and coverage are specified.

| Protein name | Score | Mass | Sign. Sequences | Coverage (%) |
| --- | --- | --- | --- | --- |
| MYCBP2 | 12344 | 517856 | 164 | 50.3 |
| CUL1 | 4091 | 90334 | 59 | 73.2 |
| FBXW7 | 3143 | 80583 | 42 | 71.9 |
| SKP1 | 1059 | 18817 | 12 | 95.7 |
| FBXO45 | 908 | 31183 | 13 | 43.4 |
| MYC | 789 | 49344 | 9 | 26.2 |
| NOTCH2 | 375 | 279081 | 4 | 4.3 |
| MED13L | 142 | 210603 | 1 | 2.6 |
| NEDD8 | 116 | 8555 | 2 | 30.3 |
| NOTCH1 | 97 | 274439 | 1 | 0.7 |
| ARIH1 | 76 | 31920 | 1 | 6.5 |
| RBX1 | 72 | 12722 | 1 | 13 |
| AURORA-B | 62 | 39977 | 0 | 6.3 |
| MED13 | 56 | 242728 | 0 | 0.9 |
| MTOR | 50 | 158783 | 0 | 1.1 |

**Figure EV1: Related to Figure 2.**
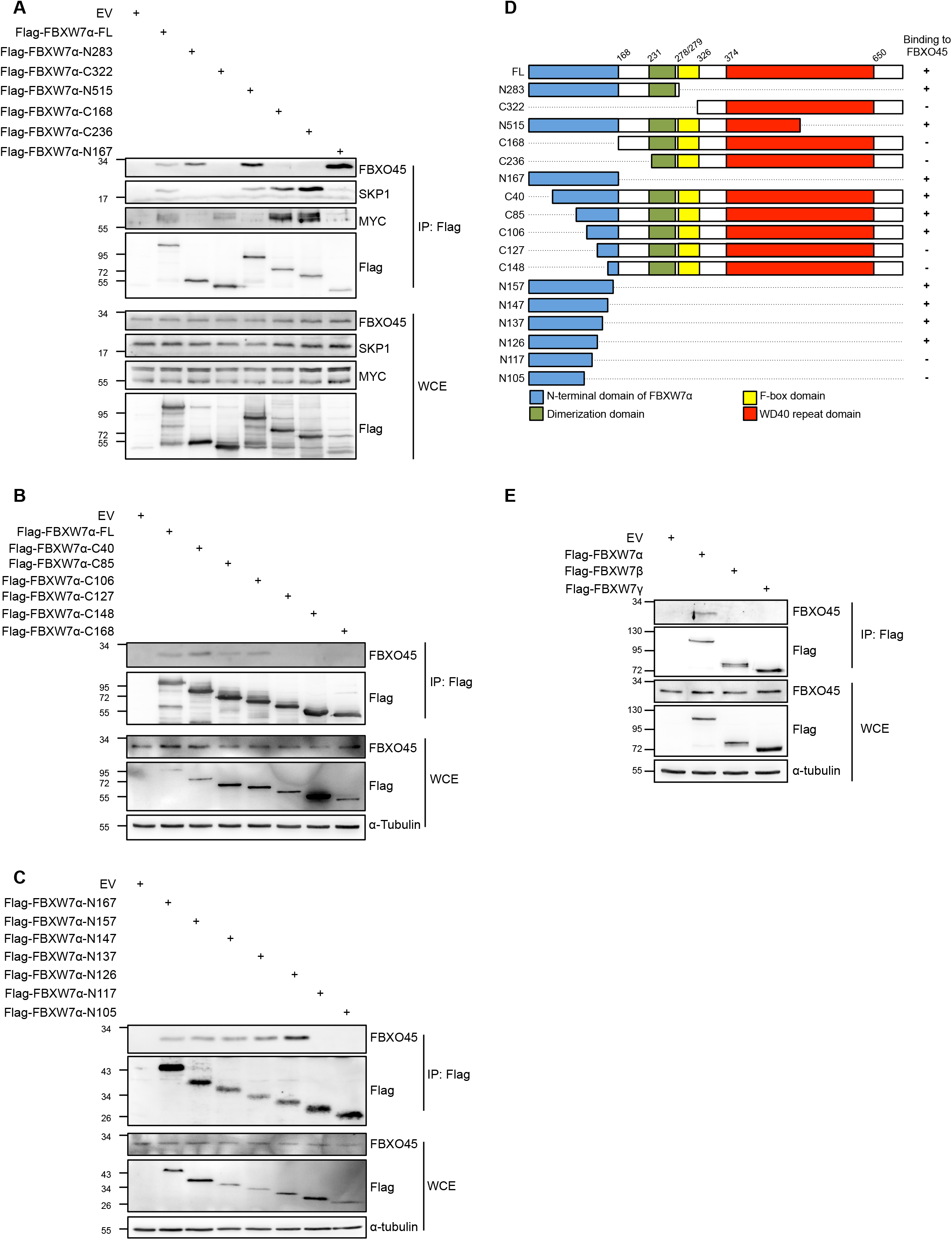
(A-C) The indicated Flag-tagged versions of FBXW7α were overexpressed in HEK-293T cells for 24 h. Cell extracts were used for immunoprecipitations with α-Flag agarose beads. (D) Overview of Flag-tagged FBXW7α fragments used in (A-C). Positions of N-terminal domain (blue), dimerization domain (green), F-box domain (yellow) and WD40 domain (red) are indicated. (E) FBXO45 specifically interacts with the α-isoform of FBXW7. Flag-FBXW7α, Flag-FBXW7β or Flag-FBXW7γ were overexpressed in HEK-293T cells for 24 h. FBXW7 isoforms were immunoprecipitated by their Flag tags. Immunoprecipitates were analyzed by Western blotting.

**Figure EV2: Related to Figure 2.**
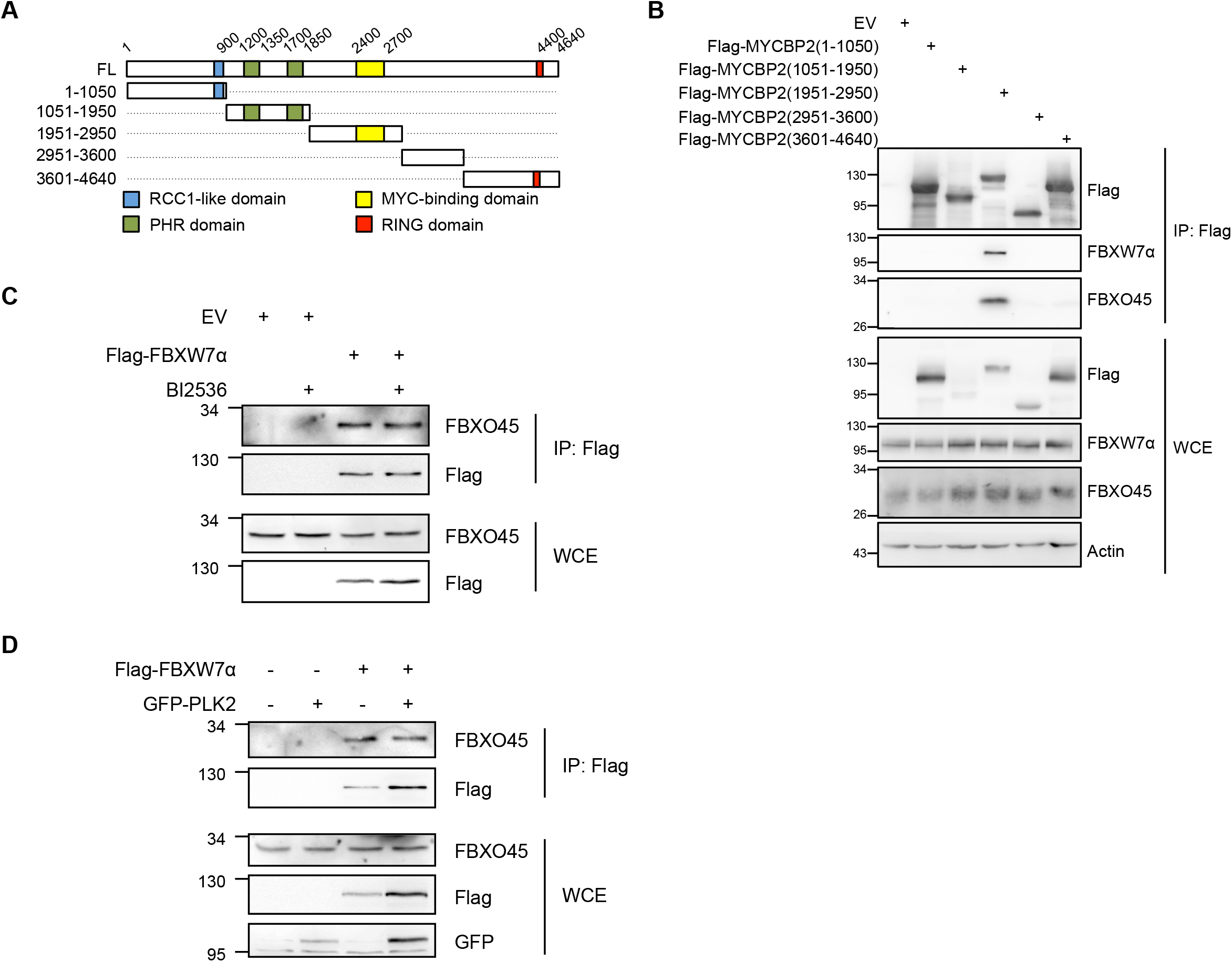
(A) Overview of MYCBP2 fragments used in (B). Positions of RCC1-like domain (blue), PHR domains (green), MYC-binding domain (yellow) and RING domain (red) are shown. (B) FBXW7α and FBXO45 interact with the central domain of MYCBP2. The indicated Flag-tagged MYCBP2 fragments were overexpressed in HEK-293T cells for 24 h. Cell extracts were used for α-Flag immunoprecipitations. (C) Flag-FBXW7α was overexpressed in HEK-293T cells for 24 h. 4 h before harvesting, the cells were treated with 100 nM BI2536. Cell extracts were used for α-Flag immunoprecipitations. (D) Flag-FBXW7α and GFP-PLK2 were co-expressed in HEK-293T cells for 24 h. Cell extracts were used for α-Flag immunoprecipitations.

**Figure EV3: Related to Figure 3.**
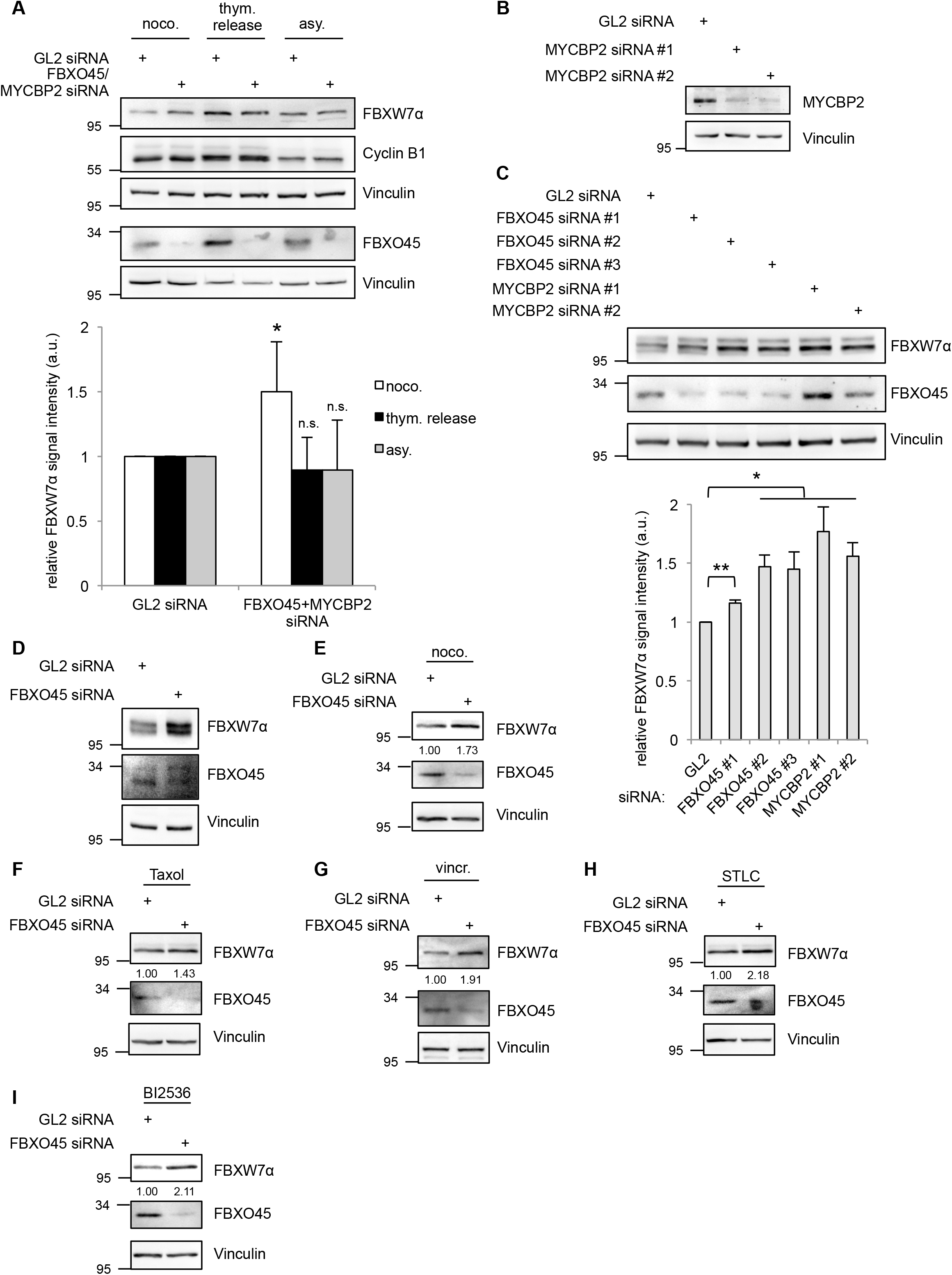
(A) HeLa cells were transfected with 30 nM FBXO45 and MYCBP2 siRNAs for 72 h. GL2 siRNA was used as a control. The cells were arrested in mitosis by nocodazole (noco.) treatment for 17 h and collected by a mitotic shake-off. Alternatively, mitotic cells were enriched after a release from a double-thymidine block (thym. release), followed by a mitotic shake-off when the cells reached the mitotic peak. Mitotic cells were compared with an asynchronous (asy.) cell population. (Top) Cell extracts were analyzed by Western blotting. (Bottom) A quantification of relative FBXW7α signal intensities is shown. Relative FBXW7α signals in the GL2 controls were set to 1. Average signal intensities and standard deviations from n=5 experiments were calculated. Statistical significance was analyzed by a two-tailed, unpaired t-test with unequal variance. * p<0.05; n.s.: not significant. (B) Characterization of MYCBP2 siRNAs used in this study. HeLa cells were transfected with 30 nM of GL2 or MYCBP2 siRNAs for 72 h. Cell extracts were analyzed by Western blotting. (C) HeLa cells were transfected with 30 nM of GL2 siRNA, three different FBXO45 siRNAs or two different MYCBP2 siRNAs for 72 h. In addition, the cells were treated with nocodazole for 17 h before they were collected by a mitotic shake-off. (Top) Cell extracts were analyzed by Western blotting. (Bottom) Relative FBXW7α signal intensities were quantified. Relative signal intensity in the GL2 control was set to 1. Average signal intensities and standard deviations from n=3 experiments were calculated. Statistical significance was analyzed by a two-tailed, unpaired t-test with unequal variance. * p<0.05; ** p<0.01. (D) U2OS cells were transfected with 30 nM of GL2 or FBXO45 siRNA for 72 h. 17 h before harvesting, they were arrested in mitosis by treatment with nocodazole. Mitotic cells were harvested by mitotic shake-off. Cell extracts were analyzed by Western blotting. (E-I) HeLa cells were transfected with 30 nM of GL2 or FBXO45 siRNA for 72 h. 17 h before harvesting, they were arrested in mitosis by treatment with nocodazole (noco., (E)), Taxol (F), vincristine (vincr., (G)), STLC (H) or BI2536 (I). Mitotic cells were harvested by mitotic shake-off. Cell extracts were prepared and analyzed by Western blotting. Relative FBXW7α signal intensities were quantified.

**Figure EV4: Related to Figure 5.**
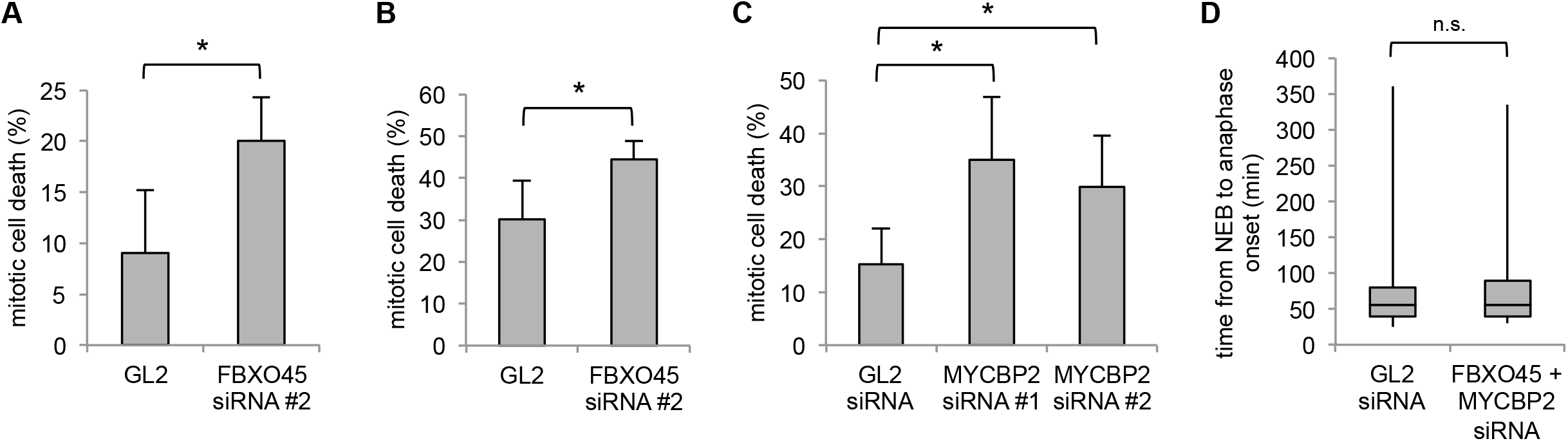
(A) U2OS cells were transfected with 30 nM of GL2 or FBXO45 siRNA for 48 h and further incubated with 1 μM Taxol. Percentages of cells undergoing mitotic cell death were quantified. Cells from n=4 independent experiments were analyzed. In each experiment, 50 cells were quantified. Statistical significance was analyzed by a two-tailed, unpaired t-test. * p<0.05. (B) U2OS cells were transfected with 30 nM of GL2 or FBXO45 siRNA for 48 h and further incubated with 1 μM vincristine. Percentages of cells undergoing mitotic cell death were quantified. Cells from n=4 independent experiments were analyzed. In each experiment, 50 cells were quantified. Statistical significance was analyzed by a two-tailed, unpaired t-test. * p<0.05. (C) U2OS cells were transfected with 30 nM of GL2 or two different MYCBP2 siRNAs for 48 h and further incubated with 250 ng/mL nocodazole. Percentages of cells undergoing mitotic cell death were quantified. Cells from n=4 independent experiments were analyzed. In each experiment, 50 cells were quantified. Statistical significance was analyzed by a two-tailed, unpaired t-test. * p<0.05. (D) HeLa H2B-RFP/GFP-α-tubulin cells were transfected with 30 nM of GL2 or FBXO45/MYCBP2 siRNAs for 72 h. They were released from a thymidine block and their mitotic progression was analyzed and quantified by live-cell imaging.

**Video S1: Related to Figure 5**

Exemplary video of a U2OS cell undergoing mitotic cell death upon prolonged mitotic arrest. Mitotic arrest was caused by treatment with nocodazole.

**Video S2: Related to Figure 5**

Exemplary video of a U2OS cell undergoing mitotic slippage upon prolonged mitotic arrest. Mitotic arrest was caused by treatment with nocodazole.

## REFERENCES

1. Hershko A, Ciechanover A (1998) The ubiquitin system. Annual review of biochemistry 67: 425–79

2. Davis RJ, Welcker M, Clurman BE (2014) Tumor suppression by the Fbw7 ubiquitin ligase: mechanisms and opportunities. Cancer cell 26: 455–64

3. Nateri AS, Riera-Sans L, Da Costa C, Behrens A (2004) The ubiquitin ligase SCFFbw7 antagonizes apoptotic JNK signaling. Science 303: 1374–8

4. Wei W, Jin J, Schlisio S, Harper JW, Kaelin WG, Jr. (2005) The v-Jun point mutation allows c-Jun to escape GSK3-dependent recognition and destruction by the Fbw7 ubiquitin ligase. Cancer cell 8: 25–33

5. Welcker M, Orian A, Jin J, Grim JE, Harper JW, Eisenman RN, Clurman BE (2004) The Fbw7 tumor suppressor regulates glycogen synthase kinase 3 phosphorylation-dependent c-Myc protein degradation. Proceedings of the National Academy of Sciences of the United States of America 101: 9085–90

6. Yada M, Hatakeyama S, Kamura T, Nishiyama M, Tsunematsu R, Imaki H, Ishida N, Okumura F, Nakayama K, Nakayama KI (2004) Phosphorylation-dependent degradation of c-Myc is mediated by the F-box protein Fbw7. The EMBO journal 23: 2116–25

7. Koepp DM, Schaefer LK, Ye X, Keyomarsi K, Chu C, Harper JW, Elledge SJ (2001) Phosphorylation-dependent ubiquitination of cyclin E by the SCFFbw7 ubiquitin ligase. Science 294: 173–7

8. Moberg KH, Bell DW, Wahrer DC, Haber DA, Hariharan IK (2001) Archipelago regulates Cyclin E levels in Drosophila and is mutated in human cancer cell lines. Nature 413: 311–6

9. Strohmaier H, Spruck CH, Kaiser P, Won KA, Sangfelt O, Reed SI (2001) Human F-box protein hCdc4 targets cyclin E for proteolysis and is mutated in a breast cancer cell line. Nature 413: 316–22

10. Hubbard EJ, Wu G, Kitajewski J, Greenwald I (1997) sel-10, a negative regulator of lin-12 activity in Caenorhabditis elegans, encodes a member of the CDC4 family of proteins. Genes & development 11: 3182–93

11. Gupta-Rossi N, Le Bail O, Gonen H, Brou C, Logeat F, Six E, Ciechanover A, Israel A (2001) Functional interaction between SEL-10, an F-box protein, and the nuclear form of activated Notch1 receptor. The Journal of biological chemistry 276: 34371–8

12. Galan JM, Peter M (1999) Ubiquitin-dependent degradation of multiple F-box proteins by an autocatalytic mechanism. Proceedings of the National Academy of Sciences of the United States of America 96: 9124–9

13. Duda DM, Olszewski JL, Tron AE, Hammel M, Lambert LJ, Waddell MB, Mittag T, DeCaprio JA, Schulman BA (2012) Structure of a glomulin-RBX1-CUL1 complex: inhibition of a RING E3 ligase through masking of its E2-binding surface. Molecular cell 47: 371–82

14. Tron AE, Arai T, Duda DM, Kuwabara H, Olszewski JL, Fujiwara Y, Bahamon BN, Signoretti S, Schulman BA, DeCaprio JA (2012) The glomuvenous malformation protein Glomulin binds Rbx1 and regulates cullin RING ligase-mediated turnover of Fbw7. Molecular cell 46: 67–78

15. Ekholm-Reed S, Goldberg MS, Schlossmacher MG, Reed SI (2013) Parkin-dependent degradation of the F-box protein Fbw7beta promotes neuronal survival in response to oxidative stress by stabilizing Mcl-1. Molecular and cellular biology 33: 3627–43

16. Cizmecioglu O, Krause A, Bahtz R, Ehret L, Malek N, Hoffmann I (2012) Plk2 regulates centriole duplication through phosphorylation-mediated degradation of Fbxw7 (human Cdc4). Journal of cell science 125: 981–92

17. Rieder CL, Maiato H (2004) Stuck in division or passing through: what happens when cells cannot satisfy the spindle assembly checkpoint. Developmental cell 7: 637–51

18. Brito DA, Rieder CL (2006) Mitotic checkpoint slippage in humans occurs via cyclin B destruction in the presence of an active checkpoint. Curr Biol 16: 1194–200

19. Frederiks CN, Lam SW, Guchelaar HJ, Boven E (2015) Genetic polymorphisms and paclitaxel-or docetaxel-induced toxicities: A systematic review. Cancer Treat Rev 41: 935–50

20. Haschka M, Karbon G, Fava LL, Villunger A (2018) Perturbing mitosis for anti-cancer therapy: is cell death the only answer? EMBO Rep 19

21. Finkin S, Aylon Y, Anzi S, Oren M, Shaulian E (2008) Fbw7 regulates the activity of endoreduplication mediators and the p53 pathway to prevent drug-induced polyploidy. Oncogene 27: 4411–21

22. Wertz IE, Kusam S, Lam C, Okamoto T, Sandoval W, Anderson DJ, Helgason E, Ernst JA, Eby M, Liu J, et al. (2011) Sensitivity to antitubulin chemotherapeutics is regulated by MCL1 and FBW7. Nature 471: 110–4

23. Grill B, Murphey RK, Borgen MA (2016) The PHR proteins: intracellular signaling hubs in neuronal development and axon degeneration. Neural Dev 11: 8

24. Po MD, Hwang C, Zhen M (2010) PHRs: bridging axon guidance, outgrowth and synapse development. Curr Opin Neurobiol 20: 100–7

25. Chen X, Sahasrabuddhe AA, Szankasi P, Chung F, Basrur V, Rangnekar VM, Pagano M, Lim MS, Elenitoba-Johnson KS (2014) Fbxo45-mediated degradation of the tumor-suppressor Par-4 regulates cancer cell survival. Cell Death Differ 21: 1535–45

26. Kugler JM, Woo JS, Oh BH, Lasko P (2010) Regulation of Drosophila vasa in vivo through paralogous cullin-RING E3 ligase specificity receptors. Molecular and cellular biology 30: 1769–82

27. Nakata K, Abrams B, Grill B, Goncharov A, Huang X, Chisholm AD, Jin Y (2005) Regulation of a DLK-1 and p38 MAP kinase pathway by the ubiquitin ligase RPM-1 is required for presynaptic development. Cell 120: 407–20

28. Xiong X, Hao Y, Sun K, Li J, Li X, Mishra B, Soppina P, Wu C, Hume RI, Collins CA (2012) The Highwire ubiquitin ligase promotes axonal degeneration by tuning levels of Nmnat protein. PLoS Biol 10: e1001440

29. Allan LA, Skowyra A, Rogers KI, Zeller D, Clarke PR (2018) Atypical APC/C-dependent degradation of Mcl-1 provides an apoptotic timer during mitotic arrest. The EMBO journal 37

30. Harley ME, Allan LA, Sanderson HS, Clarke PR (2010) Phosphorylation of Mcl-1 by CDK1-cyclin B1 initiates its Cdc20-dependent destruction during mitotic arrest. The EMBO journal 29: 2407–20

31. Kourtis N, Moubarak RS, Aranda-Orgilles B, Lui K, Aydin IT, Trimarchi T, Darvishian F, Salvaggio C, Zhong J, Bhatt K, et al. (2015) FBXW7 modulates cellular stress response and metastatic potential through HSF1 post-translational modification. Nature cell biology 17: 322–332

32. Huttlin EL, Bruckner RJ, Paulo JA, Cannon JR, Ting L, Baltier K, Colby G, Gebreab F, Gygi MP, Parzen H, et al. (2017) Architecture of the human interactome defines protein communities and disease networks. Nature 545: 505–509

33. Liao EH, Hung W, Abrams B, Zhen M (2004) An SCF-like ubiquitin ligase complex that controls presynaptic differentiation. Nature 430: 345–50

34. Welcker M, Clurman BE (2008) FBW7 ubiquitin ligase: a tumour suppressor at the crossroads of cell division, growth and differentiation. Nature reviews Cancer 8: 83–93

35. Skaar JR, Pagan JK, Pagano M (2013) Mechanisms and function of substrate recruitment by F-box proteins. Nature reviews Molecular cell biology 14: 369–81

36. Kimura T, Gotoh M, Nakamura Y, Arakawa H (2003) hCDC4b, a regulator of cyclin E, as a direct transcriptional target of p53. Cancer Sci 94: 431–6

37. Pierre SC, Hausler J, Birod K, Geisslinger G, Scholich K (2004) PAM mediates sustained inhibition of cAMP signaling by sphingosine-1-phosphate. The EMBO journal 23: 3031–40

38. Brito DA, Yang Z, Rieder CL (2008) Microtubules do not promote mitotic slippage when the spindle assembly checkpoint cannot be satisfied. The Journal of cell biology 182: 623–9

39. Gascoigne KE, Taylor SS (2008) Cancer cells display profound intra-and interline variation following prolonged exposure to antimitotic drugs. Cancer cell 14: 111–22

40. Sloss O, Topham C, Diez M, Taylor S (2016) Mcl-1 dynamics influence mitotic slippage and death in mitosis. Oncotarget 7: 5176–92

41. Cerami E, Gao J, Dogrusoz U, Gross BE, Sumer SO, Aksoy BA, Jacobsen A, Byrne CJ, Heuer ML, Larsson E, et al. (2012) The cBio cancer genomics portal: an open platform for exploring multidimensional cancer genomics data. Cancer Discov 2: 401–4

42. Gao J, Aksoy BA, Dogrusoz U, Dresdner G, Gross B, Sumer SO, Sun Y, Jacobsen A, Sinha R, Larsson E, et al. (2013) Integrative analysis of complex cancer genomics and clinical profiles using the cBioPortal. Sci Signal 6: pl1

43. Pao KC, Wood NT, Knebel A, Rafie K, Stanley M, Mabbitt PD, Sundaramoorthy R, Hofmann K, van Aalten DMF, Virdee S (2018) Activity-based E3 ligase profiling uncovers an E3 ligase with esterification activity. Nature 556: 381–385

44. Kratz AS, Richter KT, Schlosser YT, Schmitt M, Shumilov A, Delecluse HJ, Hoffmann I (2016) Fbxo28 promotes mitotic progression and regulates topoisomerase IIalpha-dependent DNA decatenation. Cell Cycle 15: 3419–3431

45. Dorr A, Pierre S, Zhang DD, Henke M, Holland S, Scholich K (2015) MYCBP2 Is a Guanosine Exchange Factor for Ran Protein and Determines Its Localization in Neurons of Dorsal Root Ganglia. The Journal of biological chemistry 290: 25620–35

46. Hoffmann I, Draetta G, Karsenti E (1994) Activation of the phosphatase activity of human cdc25A by a cdk2-cyclin E dependent phosphorylation at the G1/S transition. The EMBO journal 13: 4302–10

